# Local drug delivery to the entire cochlea without breaching its boundaries

**DOI:** 10.1101/838508

**Authors:** Andrei N. Lukashkin, Ildar I. Sadreev, Natalia Zakharova, Ian J. Russell, Yury M. Yarin

## Abstract

**SUMMARY:** The mammalian cochlea is one of the least accessible organs for drug delivery. Systemic administration of many drugs is severely limited by the blood-labyrinth barrier. Local intratympanic administration into the middle ear would be a preferable option in this case and the only option for many old and newly emerging classes of drugs but it leads to base-to-apex drug concentration gradients that are orders of magnitude and well outside the therapeutic windows. Here we present an efficient, quick, reliable and simple method of cochlear pumping, through large amplitude, low-frequency reciprocal oscillations of the stapes and round window that can consistently and uniformly deliver drugs along the entire length of the intact cochlea within minutes without disrupting cochlear boundaries. The method should facilitate novel ways of approaching the treatment of inner ear disorders since we have overcome the challenge of delivering of therapeutics along the entire length of the cochlea.

**GRAPHICAL ABSTRACT:** 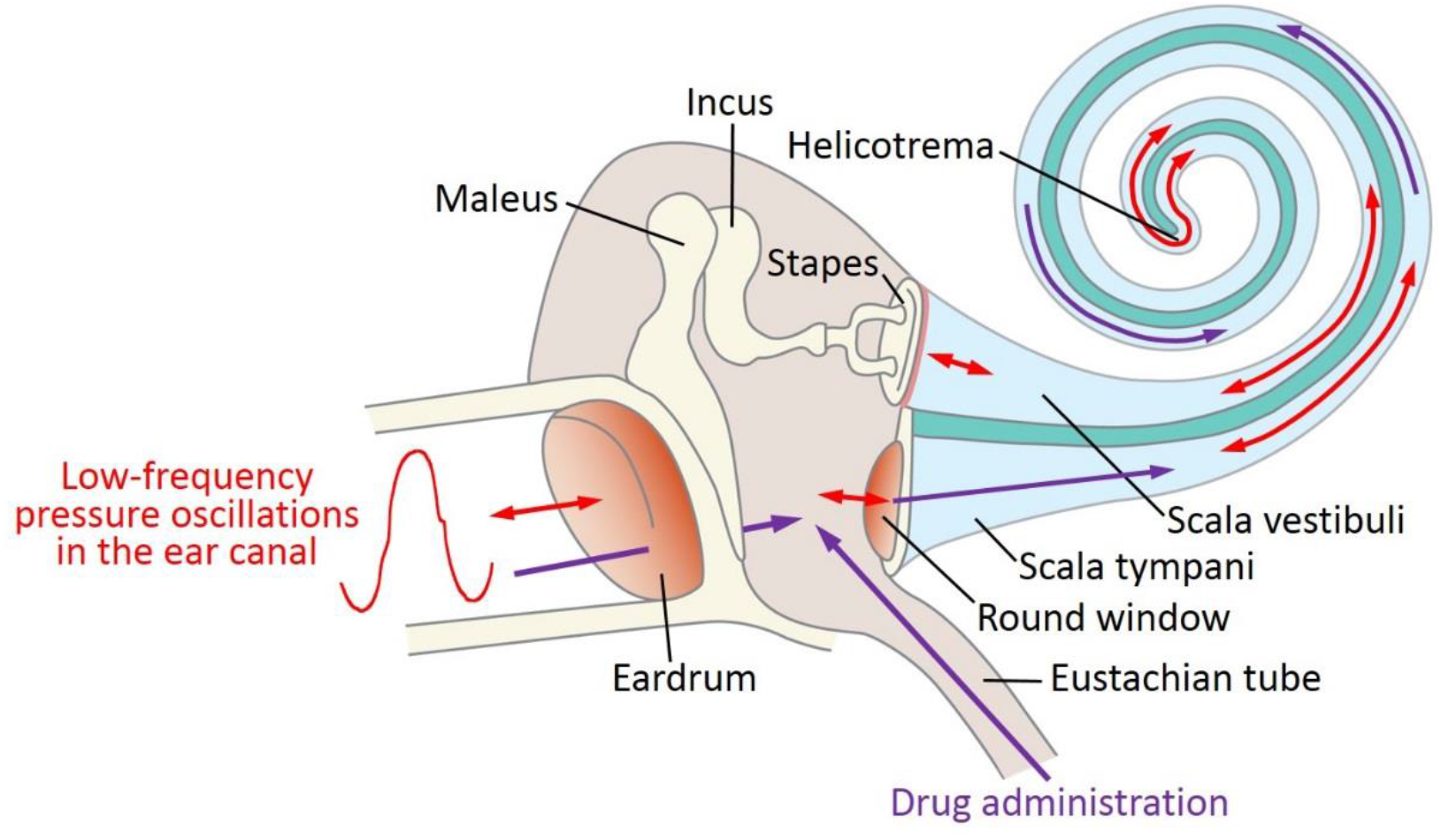

## INTRODUCTION

Reliable, efficient and uniform drug delivery to the cochlea remains an unsolved challenge and is a major barrier to the prevention or treatment of inner ear disorders. The mammalian cochlea is one of the least accessible organs for drug delivery (Salt and Plontke, 2009; Rivera et al., 2012; El Kechai et al., 2015; Hao and Li, 2019). Systemic administration of many drugs, notably the most frequently used corticosteroids and aminoglycoside antibiotics, is severely limited by the blood-labyrinth barrier (Salt and Hirose, 2018). Direct injection into the cochlea is limited by the requirement for surgery for access and does not guarantee uniform drug delivery along the cochlea.

Several potential therapeutic compounds to treat inner ear disorders are under clinical investigation. This comprises old and newly emerging classes of drugs and therapies including corticosteroids, local anaesthetics, antioxidants, apoptosis inhibitors, neurotransmitters and their antagonists, monoclonal antibodies, growth factors, signalling pathway regulators and genetic material (see Devare et al., 2018; Hao and Li, 2019). A recent review identified 43 biotech companies currently pursuing experimental compounds for inner ear therapy (Schilder et al., 2019). All such efforts are, however, restricted by our inability to reliably deliver such compounds into the cochlea.

Intratympanic administration of drugs (Schuknecht, 1956) relies on their remaining in contact with the round window (RW) (a membranous opening in the bony wall of the cochlear into the middle ear) long enough to allow their diffusion into the perilymph of the scala tympani (ST). The ability of drugs to pass through the RW does not, however, guarantee their effective distribution along the cochlear spiral. Drug distribution in the ST is limited by the low flow rate of perilymph within the cochlea and by cochlear geometry. The longitudinal flow of perilymph in the cochlea has been shown to be relatively slow, if present at all (Ohyama et al., 1988), and drug distribution in the perilymph is dominated by passive diffusion. Passive diffusion along the ST is, however, constrained because the cochlea is a relatively long and narrow tube with a cochlear cross-section that decreases gradually from the RW at the base to the apex.

Direct measurements of the distribution of marker ions and contrasting agents (Saijo and Kimura, 1984; Salt and Ma, 2001; Haghpanahi et al., 2013), corticosteroids (Hargunani et al., 2006; Plontke et al., 2008; Grewal et al., 2013; Creber et al., 2018) and antibiotics (Imamura and Adams, 2003; Mynatt et al., 2006; Plontke et al., 2007) or measurements of the physiological effects of drugs (Chen et al., 2005; Borkholder et al., 2010) have demonstrated that the concentration of substances applied to the RW is much higher in the cochlear base than in the apex.

A large number of methods, including intracochlear administration, cochleostomy and canalostomy, have been proposed for solving the problem of uniform drug distribution along the cochlea but only two current strategies address this problem without breaching the boundaries of the intact cochlea (e.g. see El Kechai et al., 2015). The first strategy relies on retaining drugs in contact with the RW to allow drug diffusion into the cochlea apex. Notable examples of devices designed for this purpose include Microwicks, osmotic pumps etc. Hydrogel-based drug delivery systems also allow retention of therapeutics in the middle ear in contact with the RW. The problem with this strategy is that retention of drugs at the RW leads to their establishing steady-state concentration gradients along the cochlea which depend on the relationship between diffusion and clearing (Salt and Ma, 2001; Sadreev et al., 2019), but the base-to-apex gradients can still be very pronounced.

The second strategy, although relatively non-invasive to the cochlea, requires development of more complex drug formulations. The technique employs drug loaded nanoparticles, which could be used to take advantage of anatomical and cellular feature of the cochlea which enable drug uptake through routes and pathways other than the ST route (Buckiová et al., 2012; Glueckert et al., 2018). Magnetically driven, drug-loaded magnetic nanoparticle can also be actively distributed along the entire cochlea (Ramaswamy et al., 2017).

Here we demonstrate that cochlear pumping, through pressure oscillations in the ear canal at frequencies low enough to avoid damage to the cochlear sensory apparatus, can consistently and uniformly deliver drugs along the entire length of the intact cochlea within minutes without disrupting cochlear boundaries.

## RESULTS AND DISCUSSION

### Cochlear pumping at low frequencies does not cause elevation of hearing thresholds

When inaudible low-frequency air pressure oscillations are presented at the ear canal, they are transmitted to the stapes which causes back and forth cochlear fluid movements through the scala vestibuli (SV) and ST coupled via helicotrema (Graphical Abstract). The RW works as a pressure relief valve during these movements and moves in counter phase with the stapes because the cochlear bony wall and fluid are poorly compressible. The low-frequency pressure changes within the cochlea are, however, shunted by the helicotrema and do not, cause stimulation of the sensory apparatus. In fact, shunting of low-frequency perilymph flow, which occurs for example during middle ear muscle reflex, thereby preventing cochlear overstimulation, is the previse role of the helicotrema (von Békésy, 1960). The high-frequency slope of the helicotrema mechanical filter is not, however, infinitely steep (6 dB/octave in guinea pigs and 12 dB/octave in humans (Marquardt et al., 2007)) and the choice of CP frequency for drug delivery is critical to prevent cochlear damage. In our experiments, low-frequency air pressure oscillations applied to the ear canal at 4 Hz, which caused large amplitude (~ 80 μm peak-to-peak) stapes displacement, did not elevate the threshold of the compound action potential (CAP) of the auditory nerve (Figure 1A).

**FIGURE 1.**
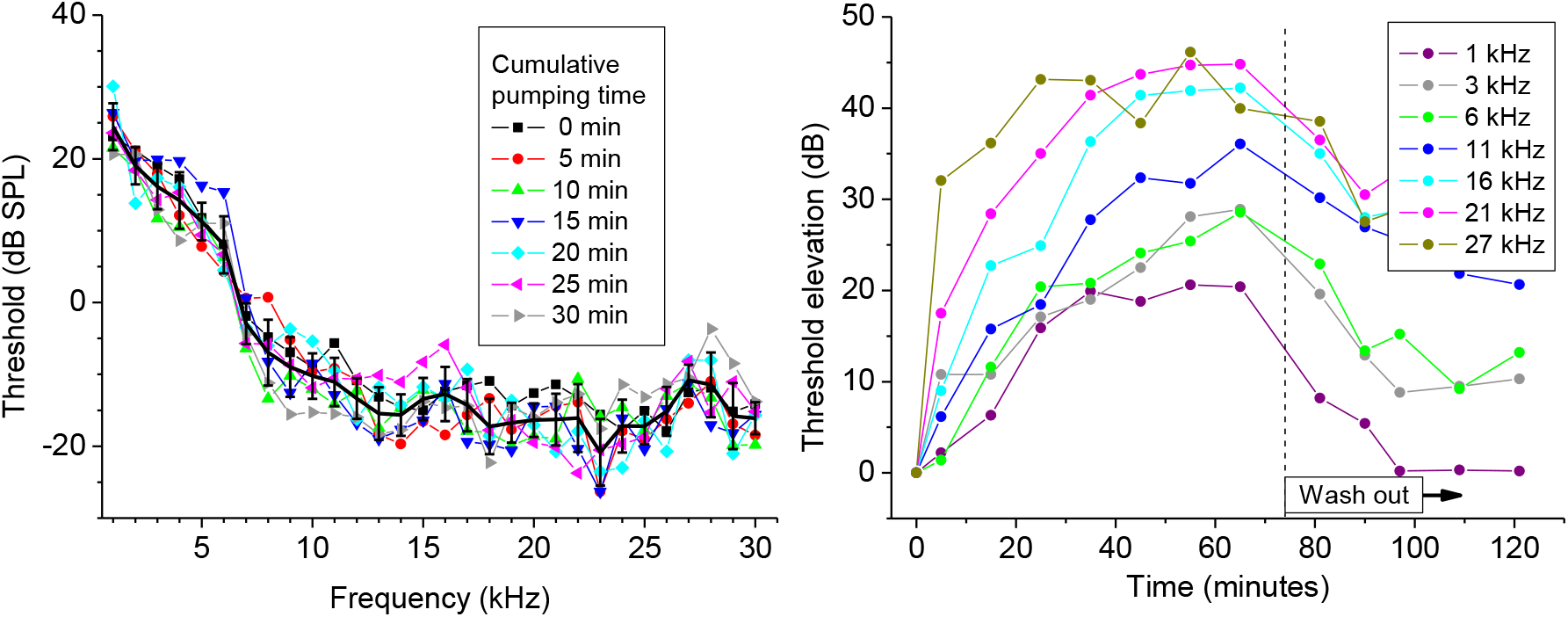
Effect of the CP on the neural responses in the absence and presence of salicylate on the RW. (A) CAP threshold curves in the presence of the CP alone. Pressure oscillations at 4 Hz that caused ~ 80 μm peak-to-peak stapes displacement were applied to the ear canal during 5 minutes before each non-zero time point plotted and the CAP thresholds were measure during the following 5-minute interval without cochlear pumping. The legend indicates the cumulative pumping time for each curve. Solid black line indicates mean ± SD for all 7 curves, each representing a different cumulative pumping time. (B) A representative examples of CAP threshold elevation when the same pumping protocol as in (A) was used after the application of 5 μl of 100 mM salicylate solution to the RW at time zero and recovery of the thresholds after washing out the salicylate. The frequency of the pure tone acoustic stimulation used for eliciting the CAP is indicated in the figure legend for each curve.

### Cochlear pumping promotes even distribution of drugs along the cochlear spiral

The ability to distribute drugs uniformly along the entire cochlea spiral using relatively large, low-frequency periodic displacement of fluid in the ST and SV (Graphical Abstract) was demonstrated in our experiments with application of salicylate to the RW. Salicylate readily passes through the RW (Borkholder et al., 2014; Sadreev et al., 2019). To monitor salicylate diffusion along an intact guinea pig cochlea *in vivo*, we utilized the suppressive effect of salicylate on cochlear amplification via block of the outer hair cell somatic motility (Russell and Schauz, 1995; Hallworth, 1997). We measured elevation of the CAP thresholds caused by salicylate at different frequencies, which, due to cochlear tonotopicity, corresponds to different distances from the RW (Greenwood, 1990).

Salicylate did not cause elevation of the CAP threshold responses for frequencies below 5 kHz, which corresponds to about 45% of the total cochlear length from the base, when it diffused through the cochlea passively (Sadreev et al., 2019). The calculated gradient of base-to-apex salicylate concentration was about 13 orders of magnitude. When, however, placement of salicylate solution on the RW was followed by cochlear pumping, i.e. by 5-minute cycles of large-amplitude (80 μm peak-to-peak), low-frequency (4 Hz) stapes movements caused by pressure oscillations in the ear canal, the CAP threshold was elevated throughout the entire 1 kHz - 30 kHz frequency range tested (Figures 1B, 2A-F, 3). This corresponds to about 75% of the total cochlear length from the base (Greenwood, 1990). Partial recovery of the CAP thresholds after washing out salicylate from the RW (Figure 1B) provided confirmation that the integrity of the sensory cells was preserved and the threshold elevation after joint salicylate application and CP was not caused by the low-frequency pressure oscillations (Figure 1A).

**FIGURE 2.**
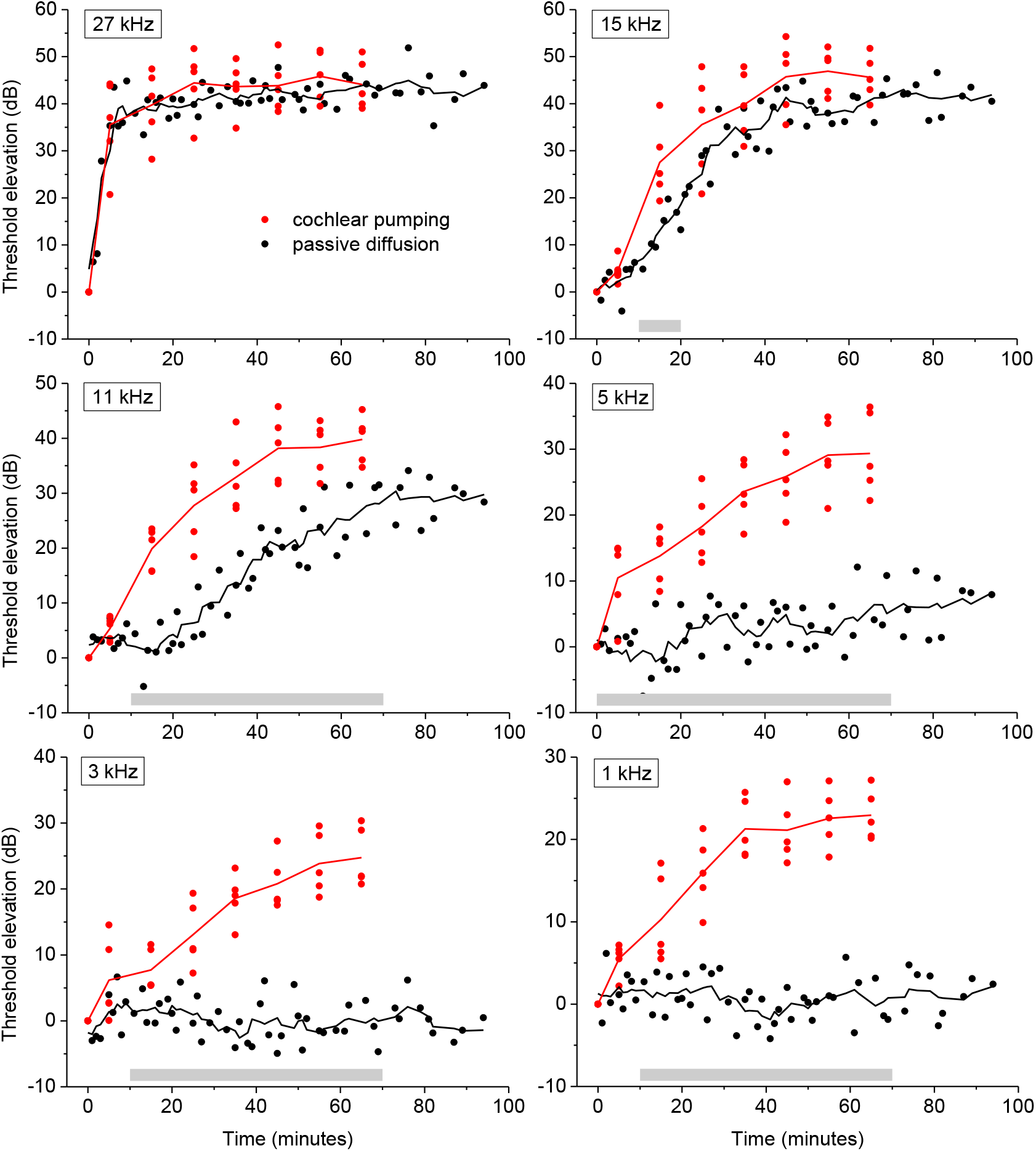
Comparison of the efficiency of the CP technique and passive diffusion in the distribution of salicylate along the cochlea. Pooled data for experiments with CP (red symbols, 5 preparations) and passive diffusion (black symbols, 5 preparations (Sadreev et al., 2019). Pressure oscillations at 4 Hz were applied to the ear canal during 5 minutes before each red non-zero time point plotted to cause large-amplitude (~ 80 μm peak-to-peak) stapes movement and the CAP thresholds were measure during the following 5-minute interval without pressure oscillation. Frequency of acoustic stimulation is indicated within each panel. 5 μl of 100 mM salicylate solution was applied to the RW at time zero. Solid red lines indicate mean CP data and solid black lines indicate 5-point running averages for passive diffusion data. Grey lines near the horizontal axis indicate statistically significant (p < 0.05, unpaired t-test) differences between data for the CP and passive diffusion within consecutive 10-minute intervals. Some of the passive diffusion data have been presented before (Sadreev et al., 2019).

**FIGURE 3.**
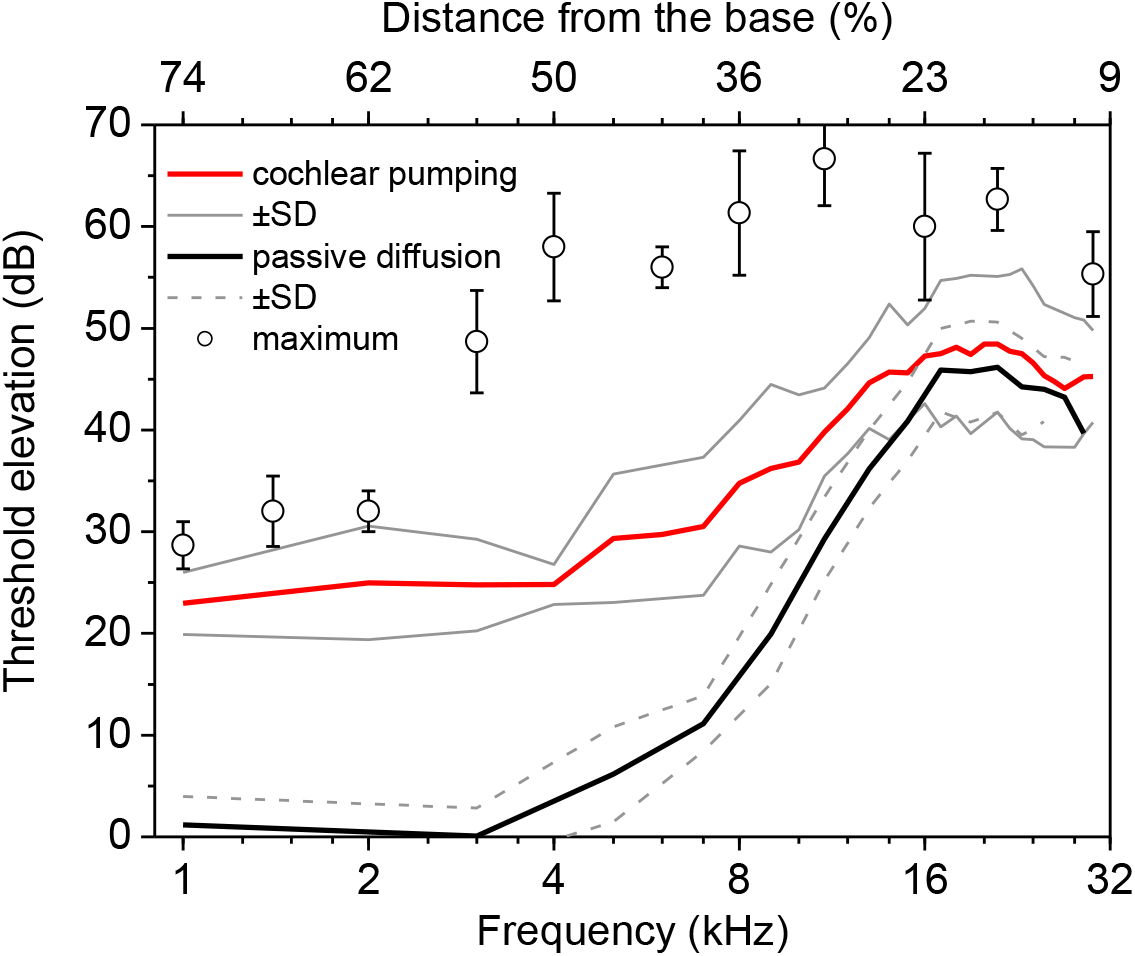
Comparison of frequency dependence of the CP and passive diffusion effects. Frequency dependence of the CAP threshold elevation after 60 minutes of salicylate application during cochlear pumping (35 minutes of the total pumping time) (mean±SD, n=5) and its comparison with the frequency dependence for passive diffusion (mean±SD, n=5). Open circles show maximal increase of the CAP thresholds after complete block of the cochlear amplifier (Sadreev et al., 2019). Data for passive diffusion have been partially presented before (Sadreev et al., 2019).

The technique had no observable influence, compared to passive diffusion, on responses to RW salicylate application for locations close to the RW at the base of the cochlea (27 kHz, Figure 2A). Cochlear pumping, however, led to more rapid threshold elevation for locations which were distal and apical to the RW, even if the maximal threshold elevations were similar for both experimental paradigms (15 kHz, Figure 2B). The threshold elevation saturated after 4-5 cycles of stimulation even for the low frequencies of the most apical locations (Figure 1B, 2E-F) where passive diffusion produced no effect. Smaller threshold elevations at low frequencies were due only to the reduced contribution of cochlear amplification to cochlear responses at these frequencies (Sadreev et al., 2019, Robles and Ruggero, 2001) and, in fact, reached almost maximal possible elevations for those frequencies (Figure 3).

A stapes displacement of 80 μm in our experiments corresponds to 1.6 mm linear displacement of the fluid in the apical parts of ST, because in guinea pigs the apical ST cross-sectional area is almost 20 times smaller (Thorne et al., 1999) than the stapes area (Sim et al., 2013) and both the cochlear bony wall and fluid are poorly compressible. Therefore, while salicylate effect in the most apical 25% of the cochlear length was not measured due to poor hearing sensitivity of guinea pigs below 1 kHz, most of the fluid in this region was replaced by fluid from more basal regions during a single cycle of cochlear pumping. Hence, estimates of salicylate distribution derived for the basal regions are valid for the most apical 25% of the cochlea.

### Tentative physical mechanisms of drug mixing and distribution during cochlear pumping

The cochlear bony wall and fluid are poorly compressible. The poor compressibility results in the fluid volume velocity along the SV and ST to be the same. Consequently, fluid linear displacement and velocity are much higher in the narrow apical parts of the scalae than at the base. Specifically, the apical ST cross-sectional area in guinea pigs is almost 20 times smaller than in the basal cochlear region (it is almost 6 times smaller in humans) (Thorne et al., 1999). This proportional increase in the fluid displacement and velocity will facilitate distribution of a drug, which originally diffuses through the RW and oval window into the cochlear base, along the entire cochlea.

The larger fluid linear velocity at the apex is still not sufficient to cause turbulent mixing of drugs. Due to the small diameter of the cochlear scalae, the fluid flow along them is dominated by fluid viscosity (i.e. it occurs at low Reynolds numbers). Thus, the fluid flow is laminar even over uneven inner surface (Cervo et al., 2013) of the scalae and turbulent mixing will not contribute to uniform drug distribution. However, the cochlear helical structure (Graphical Abstract) should lead to additional drug mixing due to the formation of Dean vortexes (e.g. Nivedita et al., 2017) and to chaotic mixing/advection, both transversal and longitudinal, observed for laminar fluid flows in helical pipes (Jones et al., 1989; Nguyen, 2011), which can be further facilitated by periodic changes of the flow direction (Ottino and Wiggins, 2004). Therefore, under specific condition of cochlear stimulation, these mechanisms may well be major factors contributing to the mixing and distribution of drugs along the ST and SV.

The tentative physical principles which govern the uniform distribution of salicylate along the cochlea are universal and should be valid for the distribution of an arbitrary substance, including nanoparticles, in the human cochlea. Salicylate was used in these initial experiments because of its well-documented physiological effect which allows estimation of the drug distribution along the intact cochlea without sampling perilymph. It was also used because it challenged the CP method. Salicylate is a difficult drug to distribute along the cochlea because it is cleared rapidly from the ST (Sadreev et al., 2019). It is anticipated that drugs, which are better retained in the ST, will be redistributed along the cochlea even more quickly and efficiently (Salt and Ma, 2001; Sadreev et al., 2019).

## ETHICS STATEMENT

All procedures involving animals were performed in accordance with UK Home Office regulations with approval from the University of Brighton Animal Welfare and Ethical Review Body.

## AUTHOR CONTRIBUTIONS

A.L., N.Z., Y.Y. and I.R. conceived and designed the study. A.L. performed the experiments. A.L. and I.S. analyzed experimental results. All authors contributed to analysis and discussion of the results. A.L. and I.R. wrote the manuscript with contribution from all authors.

## METHODS

### Animals

Animal preparation and signal generation and recording have been described elsewhere (Burwood et al., 2017). Briefly, pigmented guinea pigs of similar weight (350-360 g) and both sexes were anaesthetised with the neurolept anaesthetic technique (0.06 mg/kg body weight atropine sulphate s.c., 30 mg/kg pentobarbitone i.p., 500 μl/kg Hypnorm i.m.). Additional injections of Hypnorm were given every 40 minutes. Additional doses of pentobarbitone were administered as needed to maintain a non-reflexive state. The heart rate was monitored with a pair of skin electrodes placed on both sides of the thorax. The animals were tracheotomized and artificially respired with a mixture of O_2_/CO_2_, and their core temperature was maintained at 38°C with a heating blanket and a heated head holder. All procedures involving animals were performed in accordance with UK Home Office regulations with approval from the University of Brighton Animal Welfare and Ethical Review Body.

### Signal generation and recording

The middle ear cavity of the ear used for the measurements and salicylate application was opened to reveal the RW. Compound action potentials (CAPs) of the auditory nerve in response to pure tone stimulation were measured from the cochlear bony ridge in the proximity of the RW membrane using Teflon-coated silver wire coupled to laboratory designed and built extracellular amplifier (James Hartley). Thresholds of the N1 peak of the CAP at different frequencies which corresponds to different distances from the cochlear base (Greenwood, 1990) were estimated visually using 10 ms pure tone stimuli at a repetition rate of 10 Hz.

For acoustic stimulation sound was delivered to the tympanic membrane by a closed acoustic system comprising two Bruel and Kjaer 4134 ½” microphones for delivering tones and a single Bruel and Kjaer 4133 ½” microphone for monitoring sound pressure at the tympanum. The microphones were coupled to the ear canal via 1 cm long, 4 mm diameter tubes to a conical speculum, the 1 mm diameter opening of which was placed about 1 mm from the tympanum. The speculum was sealed in the ear canal. The closed sound system was calibrated in situ for frequencies between 1 and 50 kHz. Known sound pressure levels were expressed in dB SPL re 2×10^−5^ Pa.

All acoustic stimuli in this work were shaped with raised cosines of 0.5 ms duration at the beginning and at the end of stimulation. White noise for acoustical calibration and tone sequences for auditory stimulation were synthesised by a Data Translation 3010 board at 250 kHz and delivered to the microphones through low-pass filters (100 kHz cut-off frequency). Signals from the acoustic measuring amplifier (James Hartley) were digitised at 250 kHz using the same board and averaged in the time domain. Experimental control, data acquisition and data analysis were performed using a PC with programmes written in MATLAB (MathWorks, MA).

### Salicylate application

5 μl of 100 mM sodium salicylate solution in Hanks’ Balanced Salt Solution were placed on the RW using pipettes. The solution was removed from the RW using paper wicks to observe the wash out effect.

### Generation of pressure oscillations in the ear canal

A modified mouse ventilator MiniVent Type 845 (Hugo Sachs Elektronik, March, Germany) was used to generate oscillating air pressure in the ear canal. Output of the ventilator was connected and sealed to the closed acoustic system. The stroke frequency (4 Hz) and stroke volume (150 μl) were the same for the all experiments reported. The stroke volume was maximized to achieve maximum stapes displacement limited only by the crista stapedius.

### Recording of stapes vibrations

Stapes vibrations were recorded using a CLV-2534 laser vibrometer (Polytec GmbH, Waldbronn, Germany). The laser beam was focussed onto the stapes head. The output voltage from the vibrometer was low-pass filtered at 100 kHz, with a sensitivity of 5 mm/s/V. Stapes displacement was found by integrating the velocity responses off-line.

